# Nutrient/TOR signaling controls Mitochondrial transcription factor A (TFAM) to regulate organismal growth in *Drosophila*

**DOI:** 10.64898/2025.12.08.693086

**Authors:** Shrivani Sriskanthadevan-Pirahas, Abhishek Sharma, Shahoon Khan, Savraj S Grewal

**Affiliations:** Arnie Charbonneau Cancer Institute, Alberta Children’s Hospital Research Institute, and Department of Biochemistry and Molecular Biology Calgary, University of Calgary, Alberta T2N 4N1, Canada

**Author notes:** Authors for correspondence: SS-P and SSG.

## Abstract

Animals must adapt their growth to fluctuations in nutrient availability to ensure proper development. While nutrient-sensing tissues coordinate organismal growth through inter-organ signaling, the metabolic changes within these tissues that mediate whole-body growth control remain poorly understood. Using Drosophila larvae, we show that TOR (Target-Of-Rapamycin), a conserved nutrient-sensing kinase, controls developmental growth through regulation of the mitochondrial genome transcription factor TFAM, which controls mitochondrial bioenergetic capacity and metabolic state. We find that nutrient/TOR signaling post-transcriptionally suppresses TFAM protein levels. Furthermore, we find that TOR regulation of TFAM in the larval fat body, a key nutrient-sensing tissue, controls developmental growth. These findings establish a molecular mechanism linking nutrient-sensing pathways to mitochondrial metabolic reprogramming, revealing how environmental nutrient availability coordinates organismal growth through tissue-specific metabolic control.

## Introduction

Mitochondria are essential bioenergetic and biosynthetic organelles that generate ATP through oxidative phosphorylation (OxPhos) and provide TCA cycle products as precursors for amino acids, lipids, and nucleotides. In recent years, there has been growing interest in how these bioenergetic and biosynthetic functions couple to cell growth and proliferation, driven largely by findings that oncogenic signaling pathways alter mitochondrial function to remodel cellular metabolism in support of cancer cell growth (DeBerardinis and Chandel, 2016; 2020; DeBerardinis et al., 2008). Importantly, this work has revealed how changes in mitochondrial metabolism can actively drive cellular behaviors such as growth, proliferation, and differentiation (Homem et al., 2015; Khacho et al., 2016; Miyazawa and Aulehla, 2018; Schell et al., 2017; Senos Demarco et al., 2019). However, while much of our understanding comes from studies in cultured cells, the mechanisms operating in vivo during animal development remain less clear. In developing organisms, growth requires coordination across multiple tissues to ensure proportional size control, raising the question of how mitochondrial reprogramming in individual tissues might coordinate whole-body growth and development.

Drosophila larvae provide an excellent model system to study how growth is coordinated across tissues during animal development (Andersen et al., 2013; Boulan et al., 2015; Grewal, 2009; 2012; Koyama and Mirth, 2018; Texada et al., 2020). Larvae undergo rapid growth, increasing their mass over 200-fold during their 4–5-day development period before undergoing metamorphosis to the pupal stage. This larval growth relies critically on dietary nutrients: under nutrient-rich conditions, growth is maximal, and larvae develop rapidly, whereas nutrient-poor conditions slow growth and delay progression to pupation (Andersen *et al*., 2013; Boulan *et al*., 2015; Grewal, 2009; 2012; Koyama and Mirth, 2018; Texada *et al*., 2020). This nutrient-dependent growth is mediated largely through the conserved endocrine insulin signaling pathway. Specific neurosecretory cells in the brain—the insulin-producing cells (IPCs)—produce insulin-like peptides (dILPs) that circulate throughout the animal and stimulate growth in all tissues through activation of the insulin receptor-PI3K-Akt pathway (Grewal, 2009; 2012).

Nutrient control of this systemic insulin signaling relies on an inter-organ communication network where peripheral tissues sense nutrient availability and relay this information to the IPCs in the brain. One key nutrient-sensing tissue is the larval fat body (analogous to mammalian liver and adipose tissue). When nutrients are abundant, the fat body produces adipokines that promote dILP expression and release, sustaining high insulin signaling and rapid growth (Agrawal et al., 2016; Delanoue et al., 2016; Koyama and Mirth, 2016; Lee et al., 2018; Rajan and Perrimon, 2012; Sano et al., 2015). However, when nutrients become limiting, fat body adipokine signaling is altered, reducing insulin output and consequently slowing growth and developmental timing.

We previously demonstrated that reprogramming of mitochondrial metabolism in the fat body was a key link between nutrient input and control of systemic insulin signaling (Sriskanthadevan-Pirahas et al., 2022). We found that dietary nutrients altered fat body mitochondrial morphology, lowering bioenergetic activity. The mitochondrial transcription factor TFAM, which transcribes the mitochondrial genome, including 13 essential electron transport chain genes (Kang et al., 2007), is a key regulator of this bioenergetic capacity. When we genetically inhibited TFAM in the fat body, this alone was sufficient to accelerate larval development through alteration of adipokine signaling to the IPCs (Sriskanthadevan-Pirahas *et al*., 2022). These findings showed how fat body mitochondrial metabolic changes can non-autonomously coordinate overall body growth, but the molecular mechanisms linking nutrient signaling pathways to mitochondrial reprogramming remained unclear.

Here we explore how nutrient sensing regulates this mitochondrial metabolic reprogramming. We show that the nutrient-sensing TOR kinase signaling pathway controls TFAM protein levels and our genetic analysis demonstrates that TOR signaling in the fat body controls larval development through this TFAM-mediated pathway. This work therefore reveals a molecular link between nutrient sensing and mitochondrial metabolic reprogramming in the control of organismal growth.

## Materials and Methods

### Drosophila stocks

Flies were raised on medium containing 150 g agar, 1600 g cornmeal, 770 g Torula yeast, 675 g sucrose, 2340 g D-glucose, 240 ml acid mixture (propionic acid/phosphoric acid) per 34 L water, and maintained at 25 °C. For the low-nutrient experiments, the normal growth media was diluted to 20% in water/agar. For all GAL4/UAS experiments, homozygous GAL4 lines were crossed to the relevant UAS line(s), and the larval progeny were analyzed. The following strains were used: *w^1118^*, *r4-GAL4 (*Bloomington Drosophila Stock Center (BDSC), *UAS-TFAM RNAi (VDRC 37819 (III))*, GD control line (60000 TK), UAS-Rheb (BDSC/VDRC X).

### Measurement of *Drosophila* developmental time

For measuring development timing to the pupal stage, newly hatched larvae were collected at 24 hrs. AEL and placed in food vials (50 larvae per vial). The number of newly formed pupae was counted twice a day until all larvae had pupated. For experiments where larval development was compared in animals grown on normal vs low nutrients, larvae were raised in normal food until 70hrs AEL. They were then removed from the food by floating in 20% sucrose, washed in PBS, and then returned to either normal or low nutrient food for the remainder of their larval development, and the number of newly formed pupae was counted twice a day until all larvae had pupated.

### Mitochondrial isolation from *Drosophila* larvae

Larvae from 96 hrs grown at normal and low nutrient conditions, AEL (50 larvae per group) were frozen on dry ice or stored −80°C freezer. Larvae were thawed on ice and lysed in 500 μL of STE (250 mM sucrose, 10 mM Tris-HCl pH 7.0 and 0.2 mM EDTA pH 8.0) + BSA buffer by homogenization. Then the lysate was spun at 600 g at 4°C for 10 minutes. Supernatant was collected and centrifuged at 7 000 g for 15 min at 4°C. Next, the supernatant was discarded, and the pellet was washed in ice-cold STE buffer and resuspended in 50 μL STE buffer. The pellet containing mitochondria was aliquoted and frozen in dry ice first and transferred to – 80°C freezer for future western blot analysis. Mitochondrial protein concentration was determined using the Dc-protein determination kit (BioRad).

### Rapamycin treatment of Drosophila S2 cells

Drosophila S2 cells were cultured at 25**°**C in Schneider’s medium (Gibco; 11720-034) supplemented with 10% fetal bovine serum (Gibco; 10082-139), 100 U/ml penicillin, and 100 U/ml streptomycin (Gibco; 15140). Cells were treated with either 20 nM Rapamycin () or DMSO (Sigma; D2650) for 2 hours. After the treatment, cells were washed twice with ice-cold PBS, frozen in dry ice first, and transferred to – 80°C freezer for future analysis. Cells were then used to isolate RNA or make protein extracts as described below.

### Preparation of Drosophila S2 cells protein extracts

*Drosophila* S2 cells were lysed with a buffer containing 20 mM Tris-HCl (pH 8.0), 137 mM NaCl, 1 mM EDTA, 25 % glycerol, 1% NP-40 and with following inhibitors 50 mM NaF, 1 mM PMSF, 1 mM DTT, 5 mM sodium ortho vanadate (Na_3_VO_4_) and Protease Inhibitor cocktail (Roche Cat. No. 04693124001) and Phosphatase inhibitor (Roche Cat. No. 04906845001) according to the manufacturer’s instruction. Protein concentrations were measured using the Bio-Rad Dc Protein Assay kit II (5000112).

### Western Blot and Antibodies

Protein lysates (15 to 30μg) were resolved by SDS–PAGE and electro transferred to a nitrocellulose membrane, subjected to Western blot analysis with specific antibodies, and visualized by chemiluminescence (enhanced ECL solution (Perkin Elmer. The primary antibodies used in this study were: anti-TFAM (1:1,000, Abcam, ab47548), anti-pS6K-Thr398 (1:1000, Cell Signalling Technology #9209), anti-VDAC1 (1:1000, Santa Cruz Biotechnology sc-390996), and anti-actin (1:1000, Santa Cruz Biotechnology, # sc-8432).

### Immunostaining

*Drosophila* larvae were inverted and fixed in 8% paraformaldehyde/PBS at room temperature for 45 mins. After blocking for 2hrs in 1%BSA in PBS/0.1% Triton-X 100, larval carcasses were incubated overnight in anti-phospho S6 antibody (1:400, gift from Aurelio Teleman)). Then, inverted larvae were washed three times with 0.1% Triton X for 5 min each and incubated with an Alexa 568 goat-anti rabbit secondary antibody (1:1000) in 5% BSA in PBS for 2 hours at room temperature. Then, larvae were washed three times with 0.1 % Triton X in PBS for 5 min each, fat bodies were dissected, and mounted using VectaShield mounting medium. Larval fat bodies were imaged using a Zeiss LSM 880 with Airyscan Fast microscope and processed using Zeiss Zen Black software.

### Quantitative RT-PCR measurements

Total RNA was extracted from larvae or Drosophila S2 cells using TRIzol according to the manufacturer’s instructions (Invitrogen; 15596-018). RNA samples isolated from the same number of larvae (control vs experimental) were DNase-treated (Ambion; 2238G) and reverse transcribed using Superscript II (Invitrogen; 100004925). The generated cDNA was used as a template to perform qRT–PCRs (ABI 7500 real-time PCR system using SyBr Green PCR mix) using gene-specific primers. PCR data were normalized to RpL32. The following primers were used: RpL32 (Rp49); forward GGCCCAAGATCGTGAAGAAG, reverse-ATTTGTGCGACAGCTTAGCATATC, TFAM; forward-TGCAACAAGTTCCCCGTGAT, reverse-GCTAGGGGCCTGACTTTGTT.

### Transmission Electron Microscopy (TEM)

TEM sample processing and imaging were done as previously described. Briefly, fat bodies were dissected from different stages (72hr, 96hr, and 120 hr AEL) of larval development and fixed with 2% paraformaldehyde and 2.5% glutaraldehyde in 0.1 M cacodylate buffer, pH 7.4, for at least 2 hours. After washing three times with the same buffer, the supernatant was discarded, and warm 1% agarose was added over the pellet. After the agarose had cooled, the embedded pellet was cut into pieces. The pieces were post-fixed in 1% osmium tetroxide in cacodylate buffer for 1 hour, dehydrated in a water/ acetone series, and embedded in Epson resin. Ultrathin sections (∼60 nm thickness) were cut in a Leica EM UC7 ultramicrotome using a diamond knife and stained with 2% aqueous uranyl acetate and Reynolds’ lead citrate. The sections were examined in a Hitachi H7650 transmission electron microscope at 80 kV. The images were acquired through an AMT 16000 CCD mounted on the microscope. TEM images were analyzed using NIH Image J software by measuring the mitochondrial cross-sectional area, cell area, and quantifying the inner membrane junctions.

### Statistical analysis

All qRT-PCR data and quantification of immunostaining data were analyzed by Student’s t-test, or Mann-Whitney U test, where appropriate. All statistical analyses and data plots were performed using Prism statistical software. Differences were considered significant when p-values were less than 0.05.

## Results

### TOR and mitochondrial activity show inverse developmental dynamics

We previously showed how changes in fat body mitochondrial morphology are linked to nutrient-dependent growth. In rich nutrient conditions, mitochondria are large with fewer cristae, leading to lower bioenergetic activity. However, when nutrients are limited, mitochondria switch morphology to display dense, tighter cristae, which increases bioenergetic capacity (Sriskanthadevan-Pirahas *et al*., 2022). We also observed that these morphological changes occurred over the course of nutrient-dependent larval growth, with fat body mitochondria having features of low bioenergetic activity midway through the larval period (72-96h AEL) when nutrient-dependent growth is at its highest rate and switching to higher bioenergetic activity at the end of the larval growth period (96h-120h AEL) when feeding ceases and nutrient-dependent growth slows.

The TOR kinase pathway is a conserved regulator of nutrient-dependent growth (Ben-Sahra and Manning, 2017; Valvezan and Manning, 2019). Its activity has been reported to vary over the larval developmental period. For example, TOR activity is high in imaginal discs during mid-to-late larval stages when growth is maximal, but then declines as growth slows toward the end of the larval period (120 h AEL) (Romero-Pozuelo et al., 2017; Strassburger et al., 2021; Zhao et al., 2024). Similarly, we observed a decrease in whole-body TOR activity as larvae approached the end of the larval period (from 96 h to 120 h AEL), as assessed by phosphorylation of S6K, a direct TOR target (Figure 1B). Based on this, we specifically examined whether TOR activity in the fat body correlates with mitochondrial changes during development. Using phospho-S6 staining (a target of S6K) as a readout of TOR activity in fat body cells, we observed high TOR activity at 72 h AEL that significantly decreased at 96 h and 120 h AEL (Figure 1C, D). This pattern showed an inverse relationship with mitochondrial bioenergetic activity (Figure 1A), suggesting that TOR signaling may suppress mitochondrial function during the growth phase. To test this hypothesis, we examined the effects of overexpressing Rheb, a small GTPase that activates TOR signaling. At 120h AEL, when TOR activity is normally low, Rheb-overexpressing cells showed altered mitochondrial morphology with enlarged mitochondria displaying fewer cristae, characteristic of low bioenergetic activity (Figure 1E). In contrast, control larvae at this stage displayed smaller mitochondria with dense cristae, indicative of high bioenergetic activity. These results demonstrate that TOR signaling and mitochondrial bioenergetic activity are inversely correlated during larval development, raising the question of how TOR might regulate mitochondrial function.

**Figure 1.**
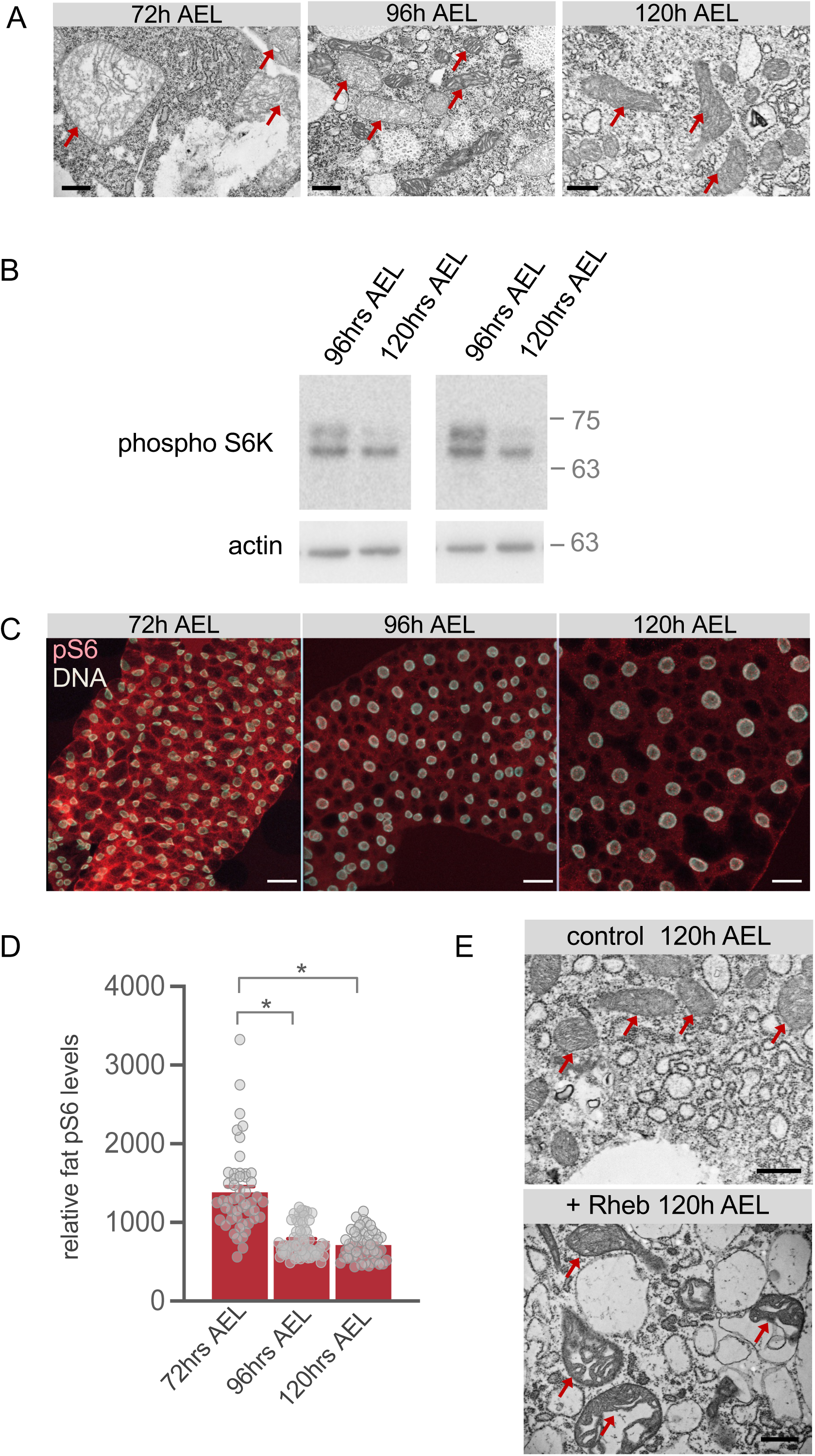
Fat-body TOR and mitochondrial activity show inverse developmental dynamics. (A) Electron micrographs of fat bodies from larvae maintained in normal food at 72 hrs, 96 hrs, or 120 hrs after egg laying (AEL). Red arrows highlight mitochondria. Scale bar represents 500 nm. (B) Western blots of whole-body samples from larvae at 96 hrs or 120 hrs AEL analyzed using anti-phospho-S6K and anti-actin antibodies. (C) Immunostaining of larval fat bodies at 72 hrs, 96 hrs, or 120 hrs after egg laying (AEL) using anti-phospho-S6 antibodies (red). DNA (white) are stained with Hoechst 33258 to visualize nuclei. Scale bar represents 40μm. (D) Quantification of phospho-S6 staining from (C). Bars represent mean +/- SEM. Symbols represent individual data points. * p < 0.05 (one-way ANOVA followed by post-hoc Tukeys test). (E) Electron micrographs of either control or Rheb overexpressing fat body cells from larvae at 120 hrs (AEL). Red arrows highlight mitochondria. Scale bar represents 500 nm.

### Nutrient availability and TOR signaling negatively regulate TFAM protein levels

To explore potential molecular mediators of TOR control of mitochondrial bioenergetics, we focused on TFAM, a nuclear-encoded transcription factor that localizes to mitochondria and transcribes the mitochondrial genome, including genes encoding essential subunits of the electron transport chain. Through this function, TFAM is a major controller of mitochondrial bioenergetic activity. Importantly, previous work in mammalian cells has shown that TFAM can be regulated by TOR signaling (Capristo et al., 2022; Chiao et al., 2016; Lerner et al., 2013; Liu et al., 2017; Morita et al., 2013). We therefore examined whether *Drosophila* TFAM might be controlled by nutrient/TOR signaling.

We first tested the effects of nutrient modulation by switching 70h AEL larvae from normal food to either continued normal food or low-nutrient food (20% dilution) for 24 hours (Figure 2A, B). TFAM mRNA levels remained unchanged between nutrient conditions. However, TFAM protein levels in isolated mitochondria were significantly elevated under low-nutrient conditions. To examine this response across a broader nutrient range, we used larvae expressing GFP-tagged TFAM and found that TFAM protein levels increased when larvae were shifted to food containing 40-60% or less of normal nutrient content, demonstrating a graded response to nutrient availability (Figure 2C).

**Figure 2.**
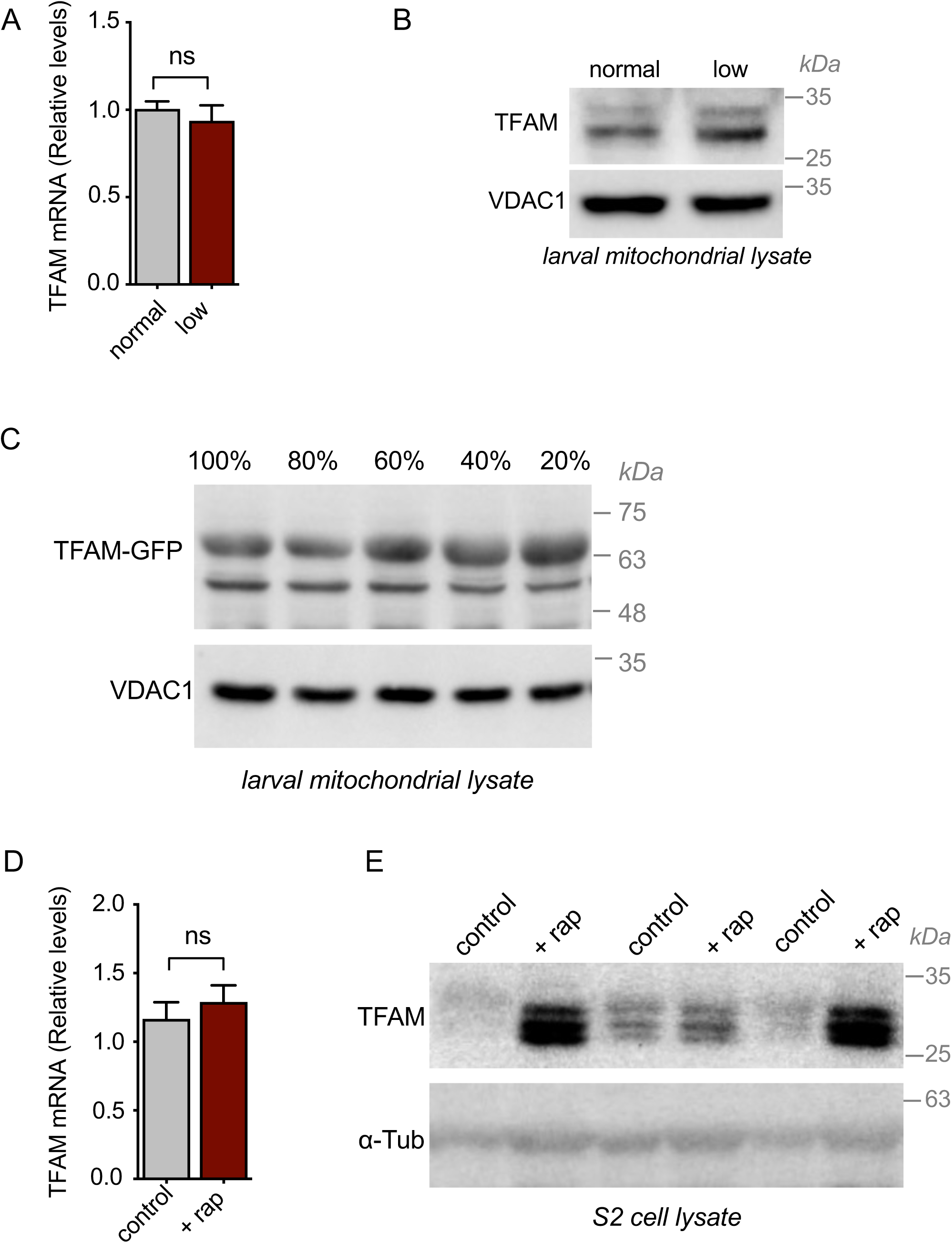
Nutrient availability and TOR signaling negatively regulate TFAM levels. (A) TFAM mRNA levels measured by qPCR from 96 hrs AEL larvae maintained on either normal food or low food for 24 hrs. Data are represented as mean ± SEM (ns, not significant unpaired t-test, n = 4 groups per condition). (B) Western blots of whole-body mitochondrial lysates from 96 hrs AEL larvae maintained on either normal food or low food for 24 hrs. Blots were analyzed using anti-TFAM and anti-VDAC1 (loading control) antibodies. (C) Western blots of whole-body mitochondrial lysates from 96 hrs AEL TFAM_GFP expressing larvae maintained on either normal food or low food for 24 hrs. Blots were analyzed using anti-GFP and anti-VDAC1 (loading control) antibodies. (D) TFAM mRNA levels measured by qPCR from control or rapamycin-treated (for 2 hrs, at 20 nM) S2 cells. Data are represented as mean ± SEM (ns, not significant unpaired t-test, n = 4 groups per condition).

To directly test whether TOR signaling controls TFAM protein levels, we used rapamycin to inhibit TOR activity in cultured *Drosophila* S2 cells (Figure 2D, E). When we examined TFAM, we found that, as with low nutrients, TFAM mRNA levels were unchanged, but TFAM protein levels were significantly increased. Together, these results demonstrate that TOR mediates nutrient control of TFAM through post-transcriptional mechanisms, likely affecting TFAM protein stability or translation.

### Negative regulation of TFAM by TOR controls developmental timing

We next examined whether TOR regulation of TFAM has functional consequences for organismal development. We previously demonstrated that lowering mitochondrial bioenergetics through TFAM knockdown accelerates development by enhancing systemic insulin signaling (Sriskanthadevan-Pirahas *et al*., 2022). Our current data show that nutrient availability and TOR signaling negatively regulate TFAM protein levels. This suggests a model in which nutrient/TOR signaling controls developmental timing through TFAM. Under nutrient-rich conditions, high TOR activity would suppress TFAM levels and mitochondrial bioenergetics, promoting rapid development. Conversely, under nutrient-poor conditions, low TOR activity would allow TFAM levels to increase, elevating mitochondrial bioenergetics and slowing development. To test this model, we examined whether TOR effects on developmental timing depend on TFAM regulation.

We first examined the effects of TOR inhibition using rapamycin feeding. Rapamycin treatment led to a delay in larval development, similar to what we previously observed with low-nutrient conditions (Figure 3A). Interestingly, when we knocked down TFAM specifically in the fat body, this was sufficient to significantly reverse the slower development caused by rapamycin treatment, suggesting that TOR promotes development through suppression of TFAM. To examine this further, we used RNAi to knockdown S6 kinase (S6K), a direct downstream target of TOR, to specifically and genetically lower TOR signaling in the fat body. Fat body S6K knockdown led to a modest delay in development that was also reversed by simultaneous TFAM knockdown (Figure 3B). These results suggest that TOR signaling in the fat body promotes developmental timing through negative regulation of TFAM.

**Figure 3.**
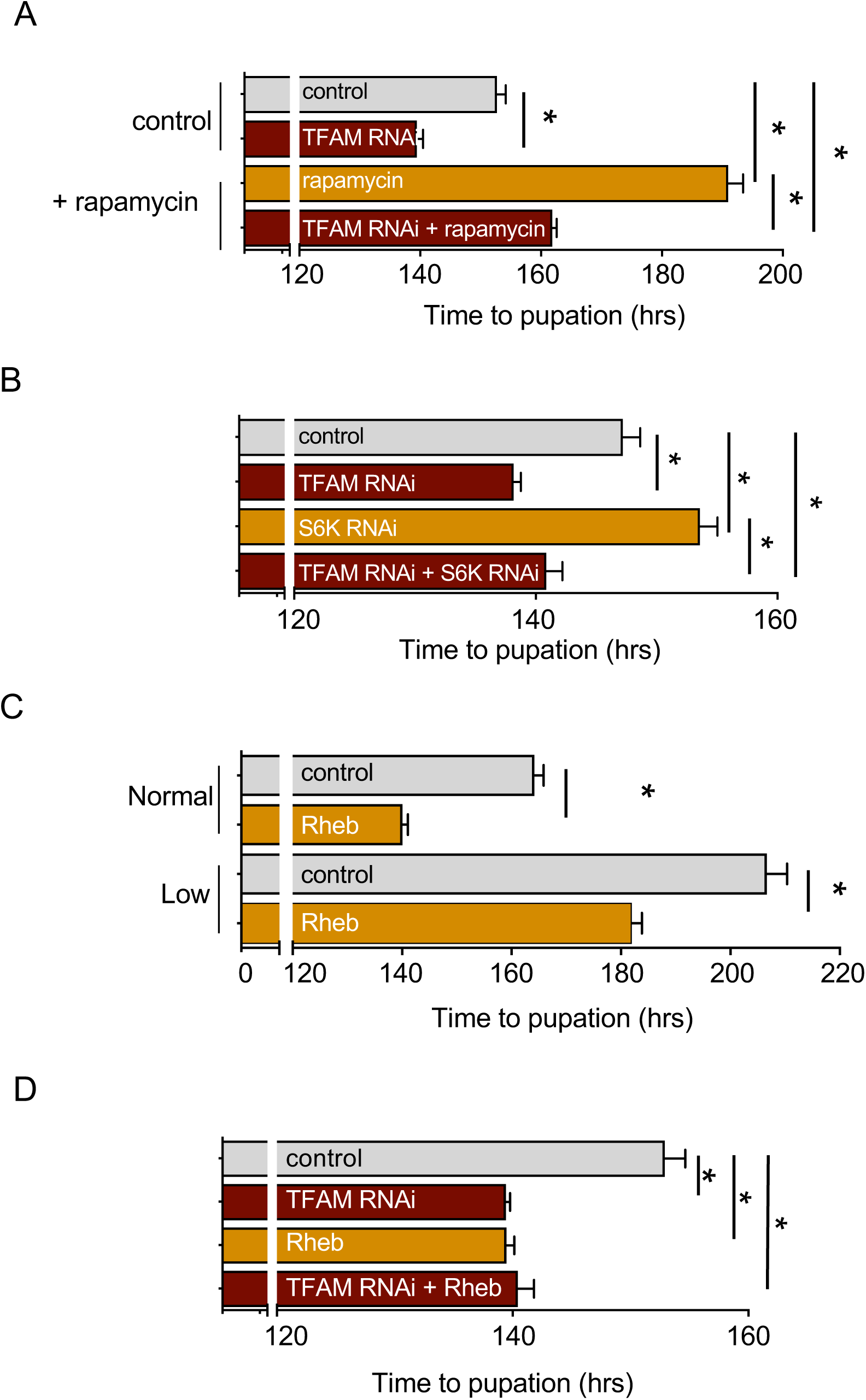
Negative regulation of TFAM by TOR controls developmental timing. (A) Time to pupation measured in control (r4-Gal4 > +) larvae vs larvae with fat body (r4-Gal4 driven) expression of TFAM RNAi grown on control or rapamycin-containing food. Data represent mean time to pupation ± SEM (*p < 0.05, ns = not significant, Mann-Whitney U test, n > 1. 50 animals per condition). (B) Time to pupation measured in control (r4 Gal4 > +) larvae vs larvae with fat body (r4-Gal4 driven) expression of TFAM RNAi, S6K RNAi or both TFAM RNAi and S6K RNAi. Data represent mean time to pupation ± SEM (*p < 0.05, ns = not significant, Mann-Whitney U test, n > 2. 125 animals per condition). (C) Time to pupation measured in control (r4-Gal4 > +) larvae vs larvae with fat body (r4-Gal4 driven) expression of UAS-Rheb grown on normal or low nutrient food. Data represent mean time to pupation ± SEM (*p < 0.05, ns = not significant, Mann-Whitney U test, n > 50 animals per condition). (D) Time to pupation measured in control (r4 Gal4 > +) larvae vs larvae with fat body (r4-Gal4 driven) expression of TFAM RNAi, Rheb or both TFAM RNAi and Rheb. Data represent mean time to pupation ± SEM (*p < 0.05, ns = not significant, Mann-Whitney U test, n > 50 animals per condition).

We next focused on examining the effects of elevated TOR signaling. Fat body-specific overexpression of Rheb was sufficient to accelerate overall development and significantly reverse the delayed development observed in low-nutrient conditions (Figure 3C). This finding suggests that modulation of fat body TOR is a main signaling link between nutrient availability and body growth. To determine whether TOR and TFAM function in the same linear pathway, we examined the genetic interaction between TFAM knockdown and Rheb overexpression. If TOR acts through TFAM, combining both manipulations should not produce additive effects. Indeed, the developmental acceleration caused by combined TFAM knockdown and Rheb overexpression was comparable to either manipulation alone (Figure 3D), demonstrating that TOR and TFAM operate in the same regulatory pathway rather than parallel pathways.

## Discussion

Our findings, together with our previous work, reveal how nutrient-dependent TOR signaling controls developmental timing through TFAM-mediated regulation of fat body mitochondrial bioenergetics (Figure 4). Under nutrient-rich conditions, high TOR activity suppresses TFAM protein levels, maintaining low mitochondrial bioenergetic activity. This metabolic state promotes systemic insulin signaling and promotes rapid organismal growth and development. Under nutrient-poor conditions, reduced TOR activity allows TFAM levels to rise, elevating mitochondrial bioenergetics, reducing insulin signaling, and slowing growth and development. Thus, TOR-mediated control of TFAM provides a molecular mechanism linking nutrient availability to tissue-specific mitochondrial reprogramming that coordinates whole-body growth.

**Figure 4.**
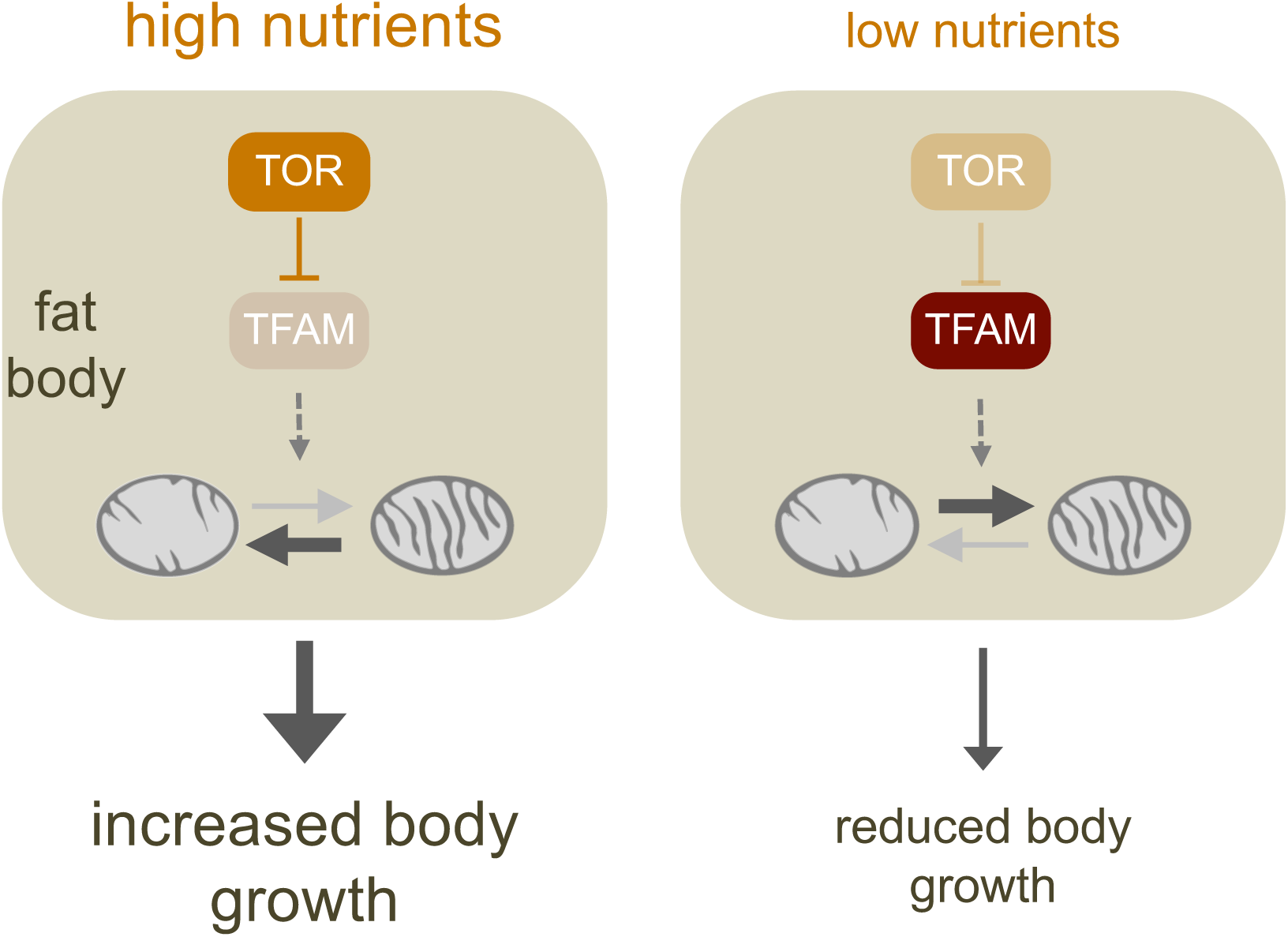
A model for Nutrient/TOR dependent control of fat body TFAM and organismal growth. Under nutrient-rich conditions, high TOR activity suppresses TFAM protein levels, maintaining low mitochondrial bioenergetic activity. This metabolic state supports rapid organismal growth and development. Under nutrient-poor conditions, reduced TOR activity allows TFAM levels to rise, elevating mitochondrial bioenergetics and slowing growth and development. Thus, TOR-mediated control of TFAM provides a molecular mechanism linking nutrient availability to tissue-specific mitochondrial reprogramming that coordinates whole-body growth

TOR control of mitochondrial metabolism has emerged as an important mechanism mediating growth control (Ben-Sahra and Manning, 2017). Extensive previous work has explored how TOR signaling regulates mitochondrial function, with several studies implicating regulation of TFAM as a key component of this control. Some of these findings are consistent with our observation that TOR negatively regulates TFAM levels. For example, rapamycin treatment increases TFAM protein in cardiac tissue and fibroblasts (Chiao *et al*., 2016; Lerner *et al*., 2013), where it contributes to the lifespan-enhancing effects of long-term rapamycin treatment.

Similarly, TOR inhibition in glioblastoma cells increases TFAM expression and promotes mitochondrial biogenesis (Ferese et al., 2020). In these mammalian systems, TOR inhibition elevates TFAM primarily through transcriptional mechanisms linked to increased autophagy. However, other reports find results opposite to ours and show that TOR can positively regulate TFAM by promoting TFAM mRNA translation through control of eIF4E-binding proteins (4EBPs), with these effects being important for mitochondrial biogenesis during cell growth, energy metabolism, and hematopoietic cell fate specification (Liu *et al*., 2017; Morita *et al*., 2013). Thus, TOR regulation of TFAM appears to be highly context-specific, operating through different mechanisms—transcriptional or translational, positive or negative—depending on cell type and physiological conditions. Our work extends this regulatory relationship to organismal development, demonstrating that post-transcriptional TOR suppression of TFAM operates in vivo to coordinate nutrient availability with whole-body growth timing through tissue-specific metabolic control.

Our results indicate that TOR post-transcriptionally suppresses TFAM protein levels in both *Drosophila* larvae and cultured S2 cells. Two potential mechanisms could explain this regulation. First, TOR might control TFAM protein stability. In mammalian cells, phosphorylation of TFAM by ERK and PKA promotes its degradation (Lu et al., 2013; Wang et al., 2014), suggesting that TOR could similarly regulate TFAM stability through phosphorylation-dependent mechanisms. Second, TOR might suppress TFAM mRNA translation. Although this appears counterintuitive given TOR’s canonical role in promoting global protein synthesis, nutrient/TOR signaling can selectively regulate translation of specific mRNAs depending on cellular context and the degree of TOR inhibition. Consistent with this, dietary restriction in *Drosophila* adults increases translation of nuclear-encoded mitochondrial genes through regulation of 4EBP, a conserved TOR effector that controls mRNA translation (Zid et al., 2009). Further biochemical studies will be needed to determine the precise mechanism by which TOR controls TFAM protein levels in *Drosophila*. Additionally, while our studies of TFAM protein levels in whole larvae and cultured cells suggest this regulation is broadly conserved across *Drosophila* cell types, direct examination of TFAM regulation specifically in larval fat body tissue will be important to confirm tissue-specific mechanisms.

Our work focused on fat body TFAM regulation in the control of growth and development. However, the fat body plays important roles in many other physiological contexts as a tissue that couples environmental conditions to systemic responses (Arrese and Soulages, 2010; Ingaramo et al., 2020; Meschi and Delanoue, 2021; Yamada et al., 2018). Similar to its role as a sensor of dietary nutrients, the fat body responds to changes in oxygen availability (Lee et al., 2019; Texada et al., 2019), pathogen exposure (Wu et al., 2012), and environmental toxins, coordinating whole-body adaptive responses that affect growth, immunity, viability, and lifespan (Arrese and Soulages, 2010; Huang and Perrimon, 2025). Interestingly, many of these environmental sensor functions involve fat body metabolic remodeling (Huang and Perrimon, 2025; Sanchez et al., 2023). This raises the possibility that the mitochondrial regulatory mechanisms we identified, particularly TOR control of TFAM, may also operate in these other contexts where the fat body must adapt its metabolic state to couple environmental changes to organismal fitness. Given that mitochondrial function is intimately linked to cellular responses to hypoxia, oxidative stress, and inflammatory signals, it will be interesting to explore whether TFAM regulation represents a broader mechanism by which the fat body integrates diverse environmental inputs to control mitochondrial metabolism.

## Acknowledgements

We thank Aurelip Teleman for the gift of the anti-phospho S6 antibody. Stocks obtained from the VDRC, the NIG-Fly Stock Centre, Kyoto, Japan and the Bloomington Drosophila Stock Center (NIH P40OD018537) were used in this study. We thank Wei-Xiang Dong for technical support with electron microscopy. This work was supported by CIHR Project Grants and a Cancer Research Society grant to S.S.G. S.K. was supported by a CIHR CGS-D award.

## References

Agrawal, N., Delanoue, R., Mauri, A., Basco, D., Pasco, M., Thorens, B., and Leopold, P. (2016). The Drosophila TNF Eiger Is an Adipokine that Acts on Insulin-Producing Cells to Mediate Nutrient Response. Cell Metab 23, 675–684. 10.1016/j.cmet.2016.03.003

Andersen, D.S., Colombani, J., and Leopold, P. (2013). Coordination of organ growth: principles and outstanding questions from the world of insects. Trends Cell Biol 23, 336–344. 10.1016/j.tcb.2013.03.005

Arrese, E.L., and Soulages, J.L. (2010). Insect fat body: energy, metabolism, and regulation. Annu Rev Entomol 55, 207–225. 10.1146/annurev-ento-112408-085356

Ben-Sahra, I., and Manning, B.D. (2017). mTORC1 signaling and the metabolic control of cell growth. Curr Opin Cell Biol 45, 72–82. 10.1016/j.ceb.2017.02.012

Boulan, L., Milan, M., and Leopold, P. (2015). The Systemic Control of Growth. Cold Spring Harb Perspect Biol 7. 10.1101/cshperspect.a019117

Capristo, M., Del Dotto, V., Tropeano, C.V., Fiorini, C., Caporali, L., La Morgia, C., Valentino, M.L., Montopoli, M., Carelli, V., and Maresca, A. (2022). Rapamycin rescues mitochondrial dysfunction in cells carrying the m.8344A > G mutation in the mitochondrial tRNA(Lys). Mol Med 28, 90. 10.1186/s10020-022-00519-z

Chiao, Y.A., Kolwicz, S.C., Basisty, N., Gagnidze, A., Zhang, J., Gu, H., Djukovic, D., Beyer, R.P., Raftery, D., MacCoss, M., et al. (2016). Rapamycin transiently induces mitochondrial remodeling to reprogram energy metabolism in old hearts. Aging 8, 314–327. 10.18632/aging.100881

DeBerardinis, R.J., and Chandel, N.S. (2016). Fundamentals of cancer metabolism. Sci Adv 2, e1600200. 10.1126/sciadv.1600200

DeBerardinis, R.J., and Chandel, N.S. (2020). We need to talk about the Warburg effect. Nat Metab 2, 127–129. 10.1038/s42255-020-0172-2

DeBerardinis, R.J., Lum, J.J., Hatzivassiliou, G., and Thompson, C.B. (2008). The biology of cancer: metabolic reprogramming fuels cell growth and proliferation. Cell Metab 7, 11–20. 10.1016/j.cmet.2007.10.002

Delanoue, R., Meschi, E., Agrawal, N., Mauri, A., Tsatskis, Y., McNeill, H., and Leopold, P. (2016). Drosophila insulin release is triggered by adipose Stunted ligand to brain Methuselah receptor. Science 353, 1553–1556. 10.1126/science.aaf8430

Ferese, R., Lenzi, P., Fulceri, F., Biagioni, F., Fabrizi, C., Gambardella, S., Familiari, P., Frati, A., Limanaqi, F., and Fornai, F. (2020). Quantitative Ultrastructural Morphometry and Gene Expression of mTOR-Related Mitochondriogenesis within Glioblastoma Cells. Int J Mol Sci 21. 10.3390/ijms21134570

Grewal, S.S. (2009). Insulin/TOR signaling in growth and homeostasis: a view from the fly world. Int J Biochem Cell Biol 41, 1006–1010. 10.1016/j.biocel.2008.10.010

Grewal, S.S. (2012). Controlling animal growth and body size - does fruit fly physiology point the way? F1000 biology reports 4, 12. 10.3410/B4-12

Homem, C.C., Repic, M., and Knoblich, J.A. (2015). Proliferation control in neural stem and progenitor cells. Nature reviews. Neuroscience 16, 647–659. 10.1038/nrn4021

Huang, K., and Perrimon, N. (2025). Metabolic command centers in Drosophila: how the fat body and oenocytes orchestrate immunity, reproduction, and aging. Curr Opin Insect Sci 73, 101459. 10.1016/j.cois.2025.101459

Ingaramo, M.C., Sanchez, J.A., Perrimon, N., and Dekanty, A. (2020). Fat Body p53 Regulates Systemic Insulin Signaling and Autophagy under Nutrient Stress via Drosophila Upd2 Repression. Cell Rep 33, 108321. 10.1016/j.celrep.2020.108321

Kang, D., Kim, S.H., and Hamasaki, N. (2007). Mitochondrial transcription factor A (TFAM): roles in maintenance of mtDNA and cellular functions. Mitochondrion 7, 39–44. 10.1016/j.mito.2006.11.017

Khacho, M., Clark, A., Svoboda, D.S., Azzi, J., MacLaurin, J.G., Meghaizel, C., Sesaki, H., Lagace, D.C., Germain, M., Harper, M.E., et al. (2016). Mitochondrial Dynamics Impacts Stem Cell Identity and Fate Decisions by Regulating a Nuclear Transcriptional Program. Cell stem cell 19, 232–247. 10.1016/j.stem.2016.04.015

Koyama, T., and Mirth, C.K. (2016). Growth-Blocking Peptides As Nutrition-Sensitive Signals for Insulin Secretion and Body Size Regulation. PLoS Biol 14, e1002392. 10.1371/journal.pbio.1002392

Koyama, T., and Mirth, C.K. (2018). Unravelling the diversity of mechanisms through which nutrition regulates body size in insects. Curr Opin Insect Sci 25, 1–8. 10.1016/j.cois.2017.11.002

Lee, B., Barretto, E.C., and Grewal, S.S. (2019). TORC1 modulation in adipose tissue is required for organismal adaptation to hypoxia in Drosophila. Nat Commun 10, 1878. 10.1038/s41467-019-09643-7

Lee, G.J., Han, G., Yun, H.M., Lim, J.J., Noh, S., Lee, J., and Hyun, S. (2018). Steroid signaling mediates nutritional regulation of juvenile body growth via IGF-binding protein in Drosophila. Proc Natl Acad Sci U S A 115, 5992–5997. 10.1073/pnas.1718834115

Lerner, C., Bitto, A., Pulliam, D., Nacarelli, T., Konigsberg, M., Van Remmen, H., Torres, C., and Sell, C. (2013). Reduced mammalian target of rapamycin activity facilitates mitochondrial retrograde signaling and increases life span in normal human fibroblasts. Aging cell 12, 966–977. 10.1111/acel.12122

Liu, X., Zhang, Y., Ni, M., Cao, H., Signer, R.A.J., Li, D., Li, M., Gu, Z., Hu, Z., Dickerson, K.E., et al. (2017). Regulation of mitochondrial biogenesis in erythropoiesis by mTORC1-mediated protein translation. Nat Cell Biol 19, 626–638. 10.1038/ncb3527

Lu, B., Lee, J., Nie, X., Li, M., Morozov, Y.I., Venkatesh, S., Bogenhagen, D.F., Temiakov, D., and Suzuki, C.K. (2013). Phosphorylation of human TFAM in mitochondria impairs DNA binding and promotes degradation by the AAA+ Lon protease. Mol Cell 49, 121–132. 10.1016/j.molcel.2012.10.023

Meschi, E., and Delanoue, R. (2021). Adipokine and fat body in flies: Connecting organs. Mol Cell Endocrinol 533, 111339. 10.1016/j.mce.2021.111339

Miyazawa, H., and Aulehla, A. (2018). Revisiting the role of metabolism during development. Development 145. 10.1242/dev.131110

Morita, M., Gravel, S.P., Chenard, V., Sikstrom, K., Zheng, L., Alain, T., Gandin, V., Avizonis, D., Arguello, M., Zakaria, C., et al. (2013). mTORC1 controls mitochondrial activity and biogenesis through 4E-BP-dependent translational regulation. Cell Metab 18, 698–711. 10.1016/j.cmet.2013.10.001

Rajan, A., and Perrimon, N. (2012). Drosophila cytokine unpaired 2 regulates physiological homeostasis by remotely controlling insulin secretion. Cell 151, 123–137. 10.1016/j.cell.2012.08.019

Romero-Pozuelo, J., Demetriades, C., Schroeder, P., and Teleman, A.A. (2017). CycD/Cdk4 and Discontinuities in Dpp Signaling Activate TORC1 in the Drosophila Wing Disc. Dev Cell 42, 376–387 e375. 10.1016/j.devcel.2017.07.019

Sanchez, J.A., Ingaramo, M.C., Gerve, M.P., Thomas, M.G., Boccaccio, G.L., and Dekanty, A. (2023). FOXO-mediated repression of Dicer1 regulates metabolism, stress resistance, and longevity in Drosophila. Proc Natl Acad Sci U S A 120, e2216539120. 10.1073/pnas.2216539120

Sano, H., Nakamura, A., Texada, M.J., Truman, J.W., Ishimoto, H., Kamikouchi, A., Nibu, Y., Kume, K., Ida, T., and Kojima, M. (2015). The Nutrient-Responsive Hormone CCHamide-2 Controls Growth by Regulating Insulin-like Peptides in the Brain of Drosophila melanogaster. PLoS Genet 11, e1005209. 10.1371/journal.pgen.1005209

Schell, J.C., Wisidagama, D.R., Bensard, C., Zhao, H., Wei, P., Tanner, J., Flores, A., Mohlman, J., Sorensen, L.K., Earl, C.S., et al. (2017). Control of intestinal stem cell function and proliferation by mitochondrial pyruvate metabolism. Nat Cell Biol 19, 1027–1036. 10.1038/ncb3593

Senos Demarco, R., Uyemura, B.S., D’Alterio, C., and Jones, D.L. (2019). Mitochondrial fusion regulates lipid homeostasis and stem cell maintenance in the Drosophila testis. Nat Cell Biol 21, 710–720. 10.1038/s41556-019-0332-3

Sriskanthadevan-Pirahas, S., Turingan, M.J., Chahal, J.S., Thorson, E., Khan, S., Tinwala, A.Q., and Grewal, S.S. (2022). Adipose mitochondrial metabolism controls body growth by modulating systemic cytokine and insulin signaling. Cell Rep 39, 110802. 10.1016/j.celrep.2022.110802

Strassburger, K., Lutz, M., Muller, S., and Teleman, A.A. (2021). Ecdysone regulates Drosophila wing disc size via a TORC1 dependent mechanism. Nat Commun 12, 6684. 10.1038/s41467-021-26780-0

Texada, M.J., Jorgensen, A.F., Christensen, C.F., Koyama, T., Malita, A., Smith, D.K., Marple, D.F.M., Danielsen, E.T., Petersen, S.K., Hansen, J.L., et al. (2019). A fat-tissue sensor couples growth to oxygen availability by remotely controlling insulin secretion. Nat Commun 10, 1955. 10.1038/s41467-019-09943-y

Texada, M.J., Koyama, T., and Rewitz, K. (2020). Regulation of Body Size and Growth Control. Genetics 216, 269–313. 10.1534/genetics.120.303095

Valvezan, A.J., and Manning, B.D. (2019). Molecular logic of mTORC1 signalling as a metabolic rheostat. Nat Metab 1, 321–333. 10.1038/s42255-019-0038-7

Wang, K.Z., Zhu, J., Dagda, R.K., Uechi, G., Cherra, S.J, 3rd., Gusdon, A.M., Balasubramani, M., and Chu, C.T. (2014). ERK-mediated phosphorylation of TFAM downregulates mitochondrial transcription: implications for Parkinson’s disease. Mitochondrion 17, 132–140. 10.1016/j.mito.2014.04.008

Wu, S.C., Liao, C.W., Pan, R.L., and Juang, J.L. (2012). Infection-induced intestinal oxidative stress triggers organ-to-organ immunological communication in Drosophila. Cell Host Microbe 11, 410–417. 10.1016/j.chom.2012.03.004

Yamada, T., Habara, O., Kubo, H., and Nishimura, T. (2018). Fat body glycogen serves as a metabolic safeguard for the maintenance of sugar levels in Drosophila. Development 145. 10.1242/dev.158865

Zhao, Y., Alexandre, C., Kelly, G., Perez-Mockus, G., and Vincent, J.-P. (2024). Growth-induced physiological hypoxia correlates with growth deceleration during normal development. bioRxiv, 2024.2006.2004.597345. 10.1101/2024.06.04.597345

Zid, B.M., Rogers, A.N., Katewa, S.D., Vargas, M.A., Kolipinski, M.C., Lu, T.A., Benzer, S., and Kapahi, P. (2009). 4E-BP extends lifespan upon dietary restriction by enhancing mitochondrial activity in Drosophila. Cell 139, 149–160. 10.1016/j.cell.2009.07.034

